# A dual mechanism underlying retroactive shifts of auditory spatial attention: dissociating target- and distractor-related modulations of alpha lateralization

**DOI:** 10.1101/2019.12.19.882324

**Authors:** Laura-Isabelle Klatt, Stephan Getzmann, Alexandra Begau, Daniel Schneider

## Abstract

Attention can be allocated to mental representations to select information from working memory. To date, it remains ambiguous whether such retroactive shifts of attention involve the inhibition of irrelevant information or the prioritization of relevant information. Investigating asymmetries in posterior alpha-band oscillations during an auditory retroactive cueing task, we aimed at differentiating those mechanisms. Participants were cued to attend two out of three sounds in an upcoming sound array. Importantly, the resulting working memory representation contained one laterally and one centrally presented item. A centrally presented retro-cue then indicated the lateral, the central, or both items as further relevant for the task (comparing the cued item(s) to a memory probe). Time-frequency analysis revealed opposing patterns of alpha lateralization depending on target eccentricity: A contralateral decrease in alpha power in *target lateral* trials indicated the involvement of target prioritization. A contralateral increase in alpha power when the central item remained relevant (*distractor lateral* trials) suggested the de-prioritization of irrelevant information. No lateralization was observed when both items remained relevant, supporting the notion that auditory alpha lateralization is restricted to situations in which spatial information is task-relevant. Altogether, the data demonstrate that retroactive attentional deployment involves excitatory and inhibitory control mechanisms.

## 1. Introduction

In everyday life, we frequently rely on selective attention in order to focus on information that is relevant while ignoring behaviorally irrelevant sensory input. Without such an attentional filter, we would be overwhelmed by the sheer abundance of sensory information. Analogously, selective attention can operate on working memory contents that are no longer physically present in the environment. Such retroactive shifts of attention are critical in order to adapt to changing task demands and allow for an efficient allocation of limited mental storage resources. The deployment of covert spatial attention to one side in mnemonic (or perceptual) space has been linked to spatially-specific modulations of alpha oscillations^1–3^. Typically, there is a relative decrease in alpha power over posterior scalp sites contralateral to the attended location, while alpha power increases contralateral to the unattended location. Based on the gating-by-inhibition framework by Jensen and Mazaheri^4^, low alpha power has been proposed to reflect a state of high excitability in the respective neural areas, whereas high alpha power reflects the functional inhibition of task-irrelevant regions.

Analogously, two mechanisms could underlie the selection of information from working memory: shifting attention within working memory may either *facilitate* or *strengthen* the relevant information, or, on the other hand, the no longer relevant contents may be *inhibited* and thereby dropped from the focus of attention within working memory. Although many studies investigating alpha lateralization interpret their findings in terms of an inhibition account, very few have been successful in actually dissociating those two mechanisms^5^. That is largely due to the fact that the majority of studies has used lateralized stimulus displays, in which targets and distractors are presented in opposite hemifields. Thus, a lateralization of alpha power in response to a shift of attention towards a left-sided target can be likewise due to a contralateral (i.e., right-hemispheric) decrease in alpha power (reflecting target prioritization) or to an ipsilateral (i.e., left-hemispheric) increase in alpha power (reflecting distractor inhibition).

Here, we aimed at distinguishing those mechanisms using an auditory working memory paradigm. In an auditory retroactive cuing task (design adapted from a previous experiment in the visual modality: Schneider et al.^6^), participants were initially cued to attend two out of three sounds in an upcoming sound array. In any case, the resulting working memory representation contained one laterally (left or right) and one centrally presented item. A retroactive cue (retro-cue) then indicated either one (i.e., selective retro-cue) or both (i.e., neutral retro-cue) of those items as further relevant for the task, which required participants to compare the cued item(s) to a centrally presented probe stimulus. Participants were instructed to indicate whether the probe stimulus was equal to the retro-cued item(s) or not. The probe stimulus could be either a new sound (i.e., a sound that never appeared in the given trial; no response), the cued sound (i.e., the sound indicated as relevant by the retro-cue; yes response), or the non-cued sound (i.e., the sound indicated as irrelevant by the retro-cue; no response). The key aspect of this design (Fig. 1) was that only either the target (i.e., the cued item) or the distractor (i.e., the non-cued item) was lateralized. This spatial arrangement of stimuli in the sound array was essential to make sure that hemispheric asymmetries in the alpha frequency-band following the retro-cue could be unambiguously linked to either the processing of the target or the distractor. The reasoning behind this design is based on the organization of afferent auditory connections and the resulting implications for hemispheric differences in processing: It is known that the contralateral projections, transmitting auditory input, are stronger and more preponderant than the ipsilateral projections^7^. Thus, the contralateral hemisphere should predominately process the attended stimulus, whereas the ipsilateral hemisphere should predominantly process the unattended stimulus. A centrally presented sound, however, should be equally represented in both hemispheres. According to this, if the selection of information from auditory working memory involves the spatially-specific inhibition of irrelevant information at a lateralized position, a contralateral increase in alpha power should be evident when the central working memory item remains relevant (i.e., when the lateral, non-cued item becomes irrelevant; *distractor lateral condition*). Here, the non-cued item is the only lateralized stimulus; hence, alpha lateralization should be unambiguously related to the processing (i.e., inhibition) of the distractor. If, however, retroactive shifts of auditory attention involve the prioritization of relevant information, a contralateral decrease in alpha power should be evident when the lateral working memory item remains relevant (i.e., when the central, non-cued item becomes irrelevant; *target lateral condition*). Again, since the cued item is the only lateralized stimulus in that case, any hemispheric asymmetry in the alpha frequency band can solely be related to the processing of the target. Obviously, those two possibilities are not mutually exclusive and may as well both contribute to successful selection of information. Yet, the experimental design allows us to distinguish the two from one another. A third neutral retro-cue condition, in which both items remained relevant, served as a control condition. A neutral cue did neither require a re-orienting of attention within working memory nor the access to the stored spatial position of the memorized sounds. Instead, once the neutral retro-cue appeared, it was sufficient to maintain the sounds’ identities. Thus, in line with previous results, showing that alpha lateralization is limited to situations in which spatial position is a task-relevant feature^8^, we did not expect an asymmetry following neutral retro-cues.

**Figure 1.**
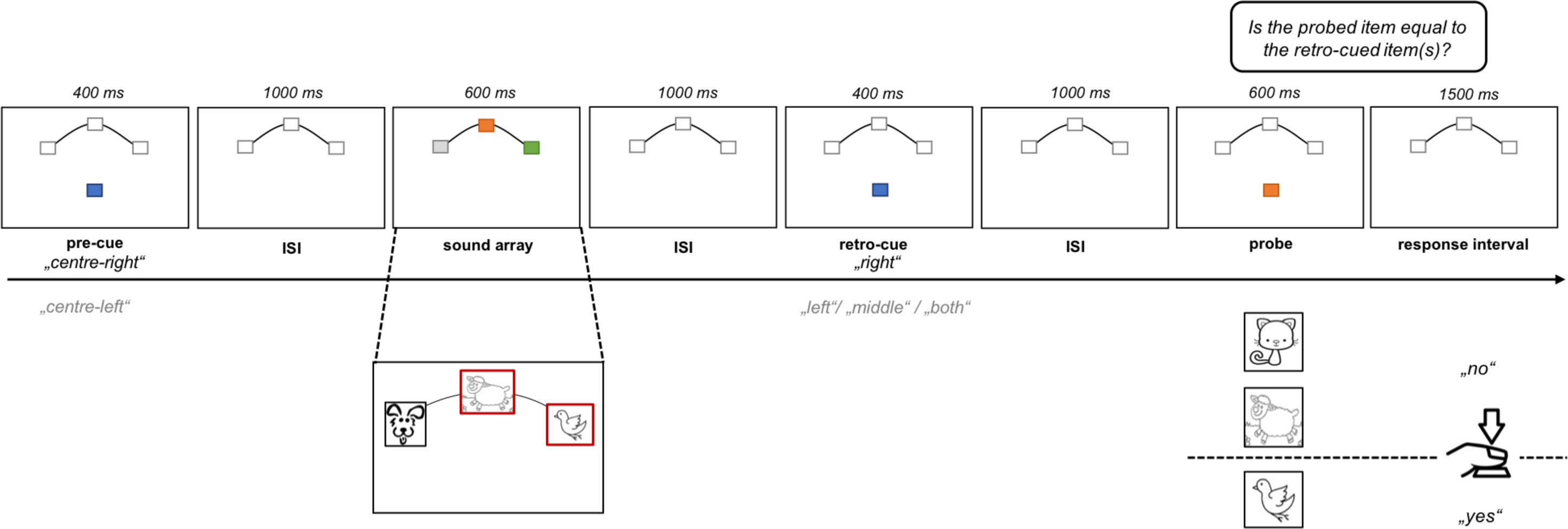
Schematic illustration of the task design. The three front loudspeakers were located at azimuthal positions of −90°, 0°, 90° in the horizontal plane. The back loudspeaker, through which pre-cue, retro-cue, and probe were presented, was located right behind the participant’s head at an approximate distance of 30 cm. Note that the depicted animal drawings exclusively illustrate that animal vocalizations were presented as sound stimuli. No visual stimuli were presented in the course of the present experiment. ISI = inter-stimulus-interval.

## 2. Results

### 2.1 Behavioral results

To investigate whether a selective retro-cue, allowing for working memory updating and a reduction in working memory load, led to an improvement in task performance compared to neutral retro-cues, we conducted a paired sample *t*-test for reaction times and a Wilcoxon signed-rank test for accuracy. Participants performed significantly faster (*t*_(19)_ = −7.93, *p* <.001, *p*_adj_ < .001, *g* = −0.60, BF > 1000) and more accurate (*z* = 2.70, *p* = .007, *p*_adj_ = .010, *U3*= .35) when only one item remained relevant (i.e., selective retro-cue trials) than when both items remained relevant (i.e., neutral retro-cue trials). In addition, to differentiate the effects of different probe types (cued, non-cued, new), we performed a one-way repeated-measures analysis of variance (rANOVA) for reaction times and a Friedman’s ANOVA for accuracy data. We expected selective retro-cue trials probing the non-cued item (which was indicated as *irrelevant* by the retro-cue) to result in slower and less accurate responses compared to trials in which the probe had never appeared in the current trial. That is, due to the interference from previously relevant information (i.e., the non-cued item), performance was expected to decline. For this comparison to work properly, neutral trials were excluded from the rANOVA, since there was no “non-cued item” when both items remained relevant. The rANOVA revealed a significant effect of probe type on reaction times (*F*_(2,38)_= 45.39, *p* <.001, *p*_adj_ = .001, *ηp*^2^ = 0.70, ε = .95) as well as on accuracy (χ^2^_(2)_ = 26.05, *p* < .001, *p*_adj_ <.001). In line with our hypotheses, post-hoc comparisons revealed that participants responded slower (*t*_(19)_ = 2.56, *p* = .019, *p*_adj_ = 0.035, *g* = 0.14, BF = 2.99) as well as less accurate (*t*_(19)_ = 8.76, *p* < .001, *p*_adj_ < .001, *g* = 1.74, BF > 1000) in trials with non-cued compared to new probes. In addition, participants responded significantly faster (*t*_(19)_ = −6.07, p < .001, *p*_adj_ <.001, *g* = −0.51, BF > 1000) as well as less accurate (*t*_(19)_ = −6.92, *p* < .001, *p*_adj_ < .001, *g* =-1.96, BF > 1000) when the cued item was probed as opposed to a new item. When contrasting trials with cued versus non-cued probes, post-hoc tests showed a significant difference in reaction times (*t*_(19)_ = −10.14, *p* < .001, *p*_adj_ < .001, *g* = −0.69, BF > 1000), but not in accuracy (*t*_(19)_ = −0.08, *p* = .934, *p*_adj_ = 1.71, *g* = −0.02, BF = 0.23). The described behavioral results are illustrated in Fig. 2 and 3.

**Figure 2.**
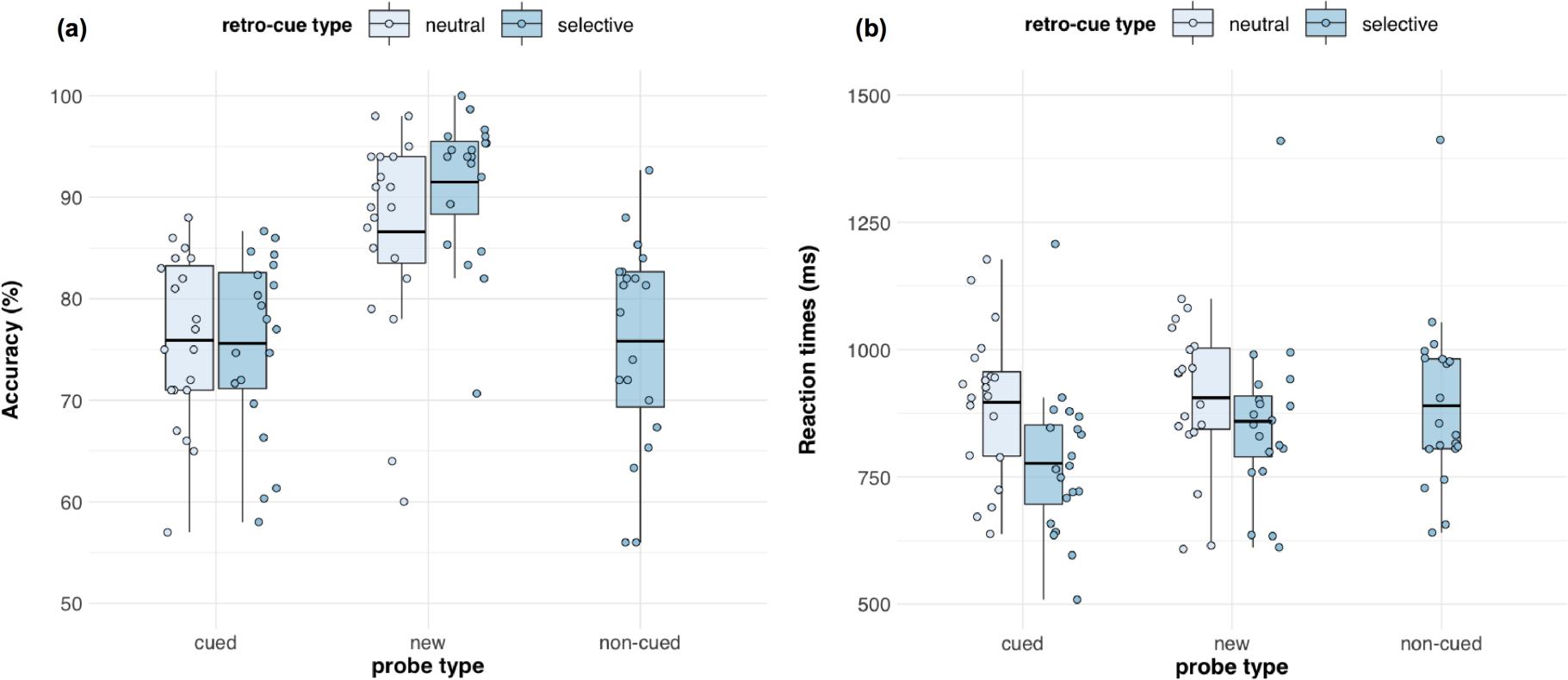
Accuracy (a) and reaction times (b), depending on retro-cue type and probe type. Boxplots illustrate the interquartile range and the mean. Whiskers extend to 1.5 times the interquartile range. The dots indicate individual participants’ mean scores per condition.

**Figure 3.**
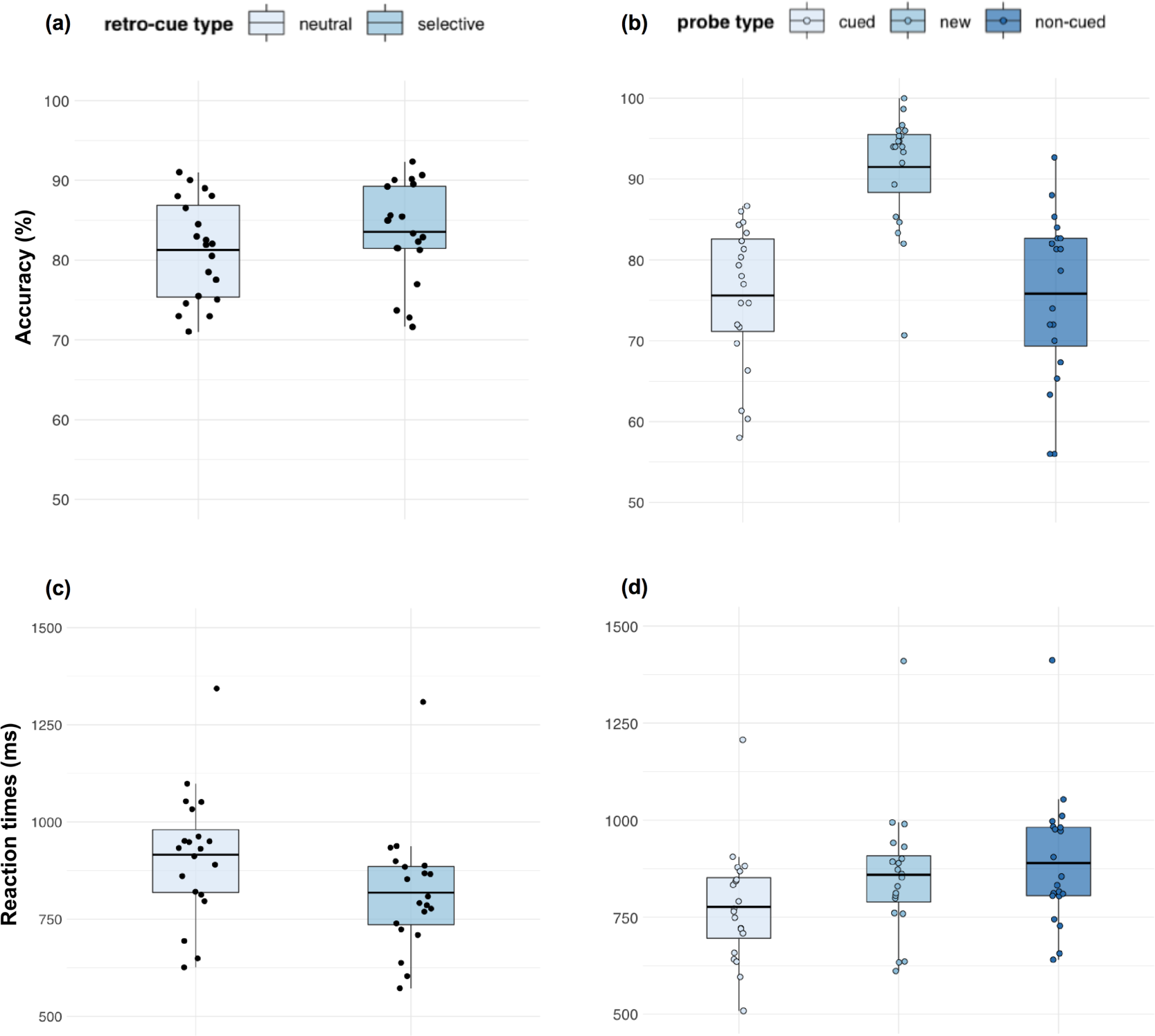
Accuracy (a, b) and reaction times (c, d), depending on the comparisons considered for statistical analyses. To test for benefits of selective retro-cues, performance in selective and neutral retro-cues was compared, using paired sample *t*-tests. Because in neutral retro-cue trials, it was not possible to probe a non-cued item, the data depicted for selective retro-cues do not include non-cued probe trials (a, c). Accuracy (a) and reaction times (d), depending on probe type, refer exclusively to selective retro-cue trials because neutral retro-cue trials did not allow for the main comparison of interest between non-cued and new items. A repeated-measured ANOVA was conducted to test for effects of probe type. For further details on statistical analyses see 5.4.1. Boxplots illustrate the interquartile range and the mean. Whiskers extend to 1.5 times the interquartile range. The dots indicate individual participants’ mean scores per condition.

### 2.2 Alpha Lateralization as a correlate of target prioritization vs. inhibition

To differentiate whether attentional selection within auditory working memory is associated with target prioritization or distractor inhibition, we contrasted the normalized difference between ipsilateral and contralateral alpha power (i.e., Alpha Lateralization Index, cf. *5.4.3*) over posterior scalp sites relative to the position of the lateralized target or distractor. As illustrated in Fig. 4, opposite patterns of lateralization were observed when comparing target lateral (Fig. 4a) and distractor lateral trials (Fig. 4b). A cluster-based permutation analysis confirmed this impression, revealing a significant difference between conditions in the alpha frequency band following the retro-cue (cluster size > 95^th^ percentile of null-distribution and *p* < .05, Fig. 4c). Follow-up analyses were performed on mean alpha power (8 – 13 Hz) in an approximate time-window derived from this comparison, ranging from 700 to 1300 ms post retro-cue onset: Paired-sample *t*-tests revealed a significant difference between neutral and target lateral trials (*t*_(19)_ = −3.45, *p* = .003, *p*_adj_ = .016, *g* = −0.98, BF = 15.46), whereas the difference between neutral trials (Fig. 4d) and distractor lateral trials failed to reach statistical significance (*t*_(19)_ = −1.17, *p* = .104, *p*_adj_ = .297, *g* = −0.52, BF =0.80). Consistent with the reported *p-*values, the BF provided strong support for the alternative hypothesis in the neutral vs. target lateral comparison, whereas it was rather inconclusive for the neutral vs. distractor lateral comparison. In order to verify that the observed hemispheric asymmetries within conditions were significantly different from zero, one-sample *t*-tests were conducted: The results suggested that there was a reliable lateralization of alpha power in the target lateral (*t*_(19)_ = 2.85, *p* = .010, *p*_adj_ = .039, *g1* = 0.64, BF = 5.07) as well as the distractor lateral condition (*t*_(19)_ = −3.58, *p* = .002, *p*_adj_ = .016, *g*_*1*_ =-0.80, BF = 20.20). With BFs greater than 3 and 10, respectively, those tests provide moderate and strong support for the alternative hypothesis (i.e., the lateralization is significantly different from zero). In contrast, there was no evidence for a lateralization of alpha power in the neutral retro-cue condition (*t*_(19)_ = −1.54, *p* = .139, *p*_adj_ = .317, *g*_*1*_ = −0.35, BF = 0.64). The corresponding BF did neither support the null nor the alternative hypothesis. Scalp topographies corresponding to the alpha lateralization indices are depicted in Fig. 4e. Related studies from the visual domain have previously raised concerns that posterior alpha power asymmetries might be confounded by lateral saccadic eye movement artifacts. See the supplementary material (Appendix A) for a control analysis, invalidating such confounding effects.

**Figure 4.**
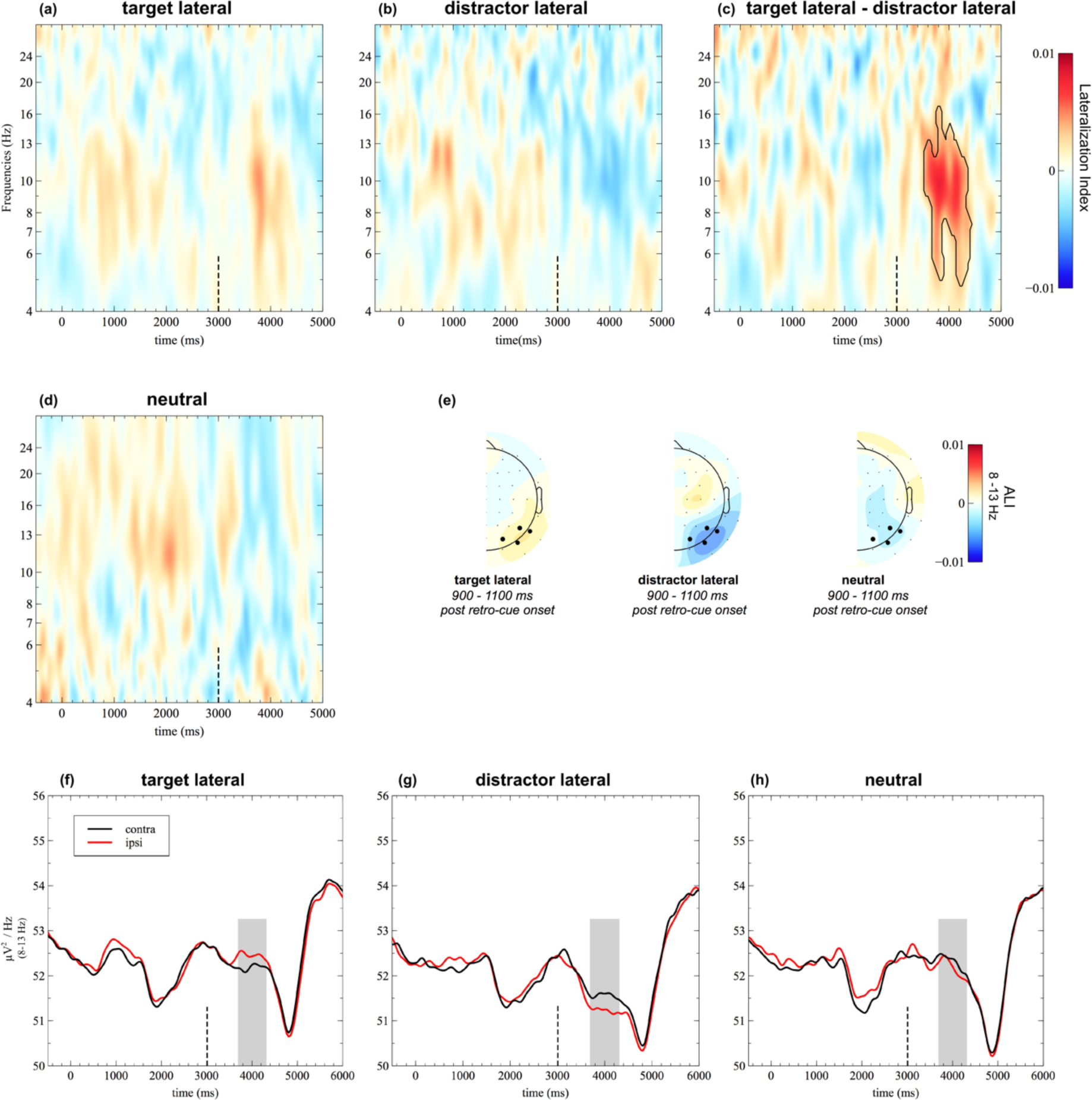
Alpha lateralization results for target lateral, distractor lateral, and neutral trials. Time-frequency plots (a, b, d) show the grand average lateralization indices at posterior electrodes (PO7/8, P7/8, P5/6, and PO3/4). The black dashed line indicates retro-cue onset. Target lateral and distractor lateral trials were contrasted, using cluster permutation statistics. The resulting significant cluster (*p* < .05 and size > 95^th^ percentile of the null distribution) is framed by a black line in the target lateral minus distractor lateral difference plot (c). Corresponding topographies are based on the normalized difference between ipsilateral and contralateral alpha power following the retro-cue (e). Line plots (f-h) illustrate the contralateral and ipsilateral portion of alpha power for the same posterior cluster of electrodes. The analysis time window is highlighted in grey. ALI = Alpha Lateralization Index.

### 2.3 Bilateral alpha power desynchronization as a measure of cognitive task demands

The line plots in Fig. 4 (f-h) clearly illustrate that there is a bilateral suppression of alpha power following retro-cue onset. Alpha desynchronization, which is linked to a state of high excitability^9^, has been associated with cognitive task demands, in particular with respect to memory and attention^10,11^ (for a review see^12^). Here, this desynchronization of alpha power appears to be more pronounced in distractor lateral trials than in target lateral or neutral trials. To statistically assess differences in alpha desynchronization between conditions, we performed a one-way repeated measures ANOVA, including the factor *retro-cue type* and bilateral, baseline-corrected alpha power (700 to 1300 ms post retro-cue onset, cf .2.2) as a dependent variable. The analysis revealed a main effect of *retro-cue type* (*F*_(2,38)_ = 6.60, *p* =.003, *ηp*^2^ = 0.26, ε = .94). Post-hoc comparisons indicated that there was a greater suppression of alpha power in distractor lateral trials compared to target lateral trials (*t*_(19)_ = 3.16, *p* = .005, *p*_adj_ = .028, *g* = 0.47, BF = 9.07) as well as compared to neutral trials (*t*_(19)_ = 2.83, *p* = .011, *p*_adj_ = .030, *g* = 0.45, BF = 4.81). In contrast, bilateral alpha suppression did not differ between target lateral and neutral trials (*t*_(19)_ = 0.07, *p* = .944, *p*_adj_ = 1.731, *g* = 0.01, BF10 = 0.23).

## 3. Discussion

The present study investigated the mechanisms underlying retroactive attentional orienting within auditory working memory. We demonstrated that signatures of target-related and distractor-related alpha lateralization can be dissociated. Specifically, by manipulating the spatial arrangement of targets and distractors such that only one of them appeared in a lateralized position, we were able to show that a retro-cue induced shift of attention towards a *lateralized target* resulted in a contralateral *decrease* in alpha power; in contrast, shifting attention to a centrally presented target and away from a *lateralized distractor* resulted in a contralateral *increase* in alpha power. The findings support the interpretation of alpha lateralization as a dual mechanism underlying the deployment of attention within working memory.

### 3.1 Retro-cue benefits: effects on accuracy and reaction times

On the behavioral level, participants clearly benefited from selective retro-cues, reducing the amount of relevant information to be maintained in working memory. While participants were initially required to maintain two sound stimuli and their respective locations, following a selective retro-cue, only one item remained relevant. This resulted in a behavioral improvement in terms of faster response times and higher accuracy compared to neutral retro-cue trials, in which both the lateral and the central item remained relevant. A large body of literature has investigated the mechanisms underlying such “*retro-cue benefits*” (for a review, see ^13^), showing that focusing attention within working memory results in the *strengthening* of attended working memory representations^14,15^ and the *removal* of non-cued information from working memory^3,16^, thereby releasing resources for further processing^17^. When looking specifically at selective retro-cue trials, we found increased response times and decreased accuracy in non-cued compared to new probe trials, suggesting a proactive interference effect in non-cued probe trials. It should be noted that although non-cued items were probed in a subset of trials, the retro-cue was always 100% valid. That is, when participants were asked to judge whether the probe item matched the cued item, it made no difference whether the probe was a new item or the previously non-cued item. As long as they remembered the (validly) cued item, they would always be able to respond correctly. Hence, the question emerges, if the non-cued item is *removed* from working memory, why does it still cause interference when presented as a probe item?

First of all, the observed interference effect is in line with retro-cue studies incorporating invalidly cued trials, which consistently suggest that non-cued items are not irreversibly lost. Those studies reliably report a decline in performance when the non-cued item was invalidly probed; however, performance never dropped to chance level^18,19^. In addition, double-cueing paradigms suggest that the information that was initially dropped from working memory (in response to a first cue) can be restored to some extent by a second retro-cue^14,20^. This is also consistent with the recently proposed framework of “activity-silent” working memory^21^, which is based on an increasing number of studies suggesting that persistent neural firing is not a necessary prerequisite for successful short-term memory maintenance. Using multivariate pattern analysis, it could be convincingly demonstrated that the active neural signature representing currently maintained information disappears once they become temporarily irrelevant. Critically, their neural signature is restored once those items are again cued to be relevant for the task^22,23^. Accordingly, these studies suggest that non-cued items are *removed from the focus of attention* and from a state of persistent neural activity, but are still available for reactivation by re-storing their neural signature. Consistently, non-cued probe items in the present study, which resulted in slower reaction times and lower accuracy, might have been (temporally) removed from the focus of attention but sequentially reactivated by the current retrieval context (i.e., the probe)^22^. Alternative explanations for such an “imperfect removal” may likewise apply, which will be further discussed below in the context of the obtained time-frequency results (cf. 3.2).

### 3.2 Alpha lateralization as a proxy of target prioritization and distractor inhibition?

On a neural level, shifts of attention in response to the retro-cue were reflected by a hemispheric lateralization of posterior alpha lateralization. Broadly speaking, this is consistent with a growing body of literature showing that attentional modulation of alpha oscillations are comparable for shifts of attention within perceptual space and within working memory^2,8,24^. Given that it is relatively undisputed that alpha lateralization tracks the attended location within working memory^25–27^, the observed spatially-specific modulations clearly indicate a re-orienting of spatial attention towards relevant items (and away from irrelevant items). Consequentially, the relevant item is moved into a prioritized state, commonly referred to as the focus of attention^28^, which renders the selected information more accessible for future cognitive operations. In line with that, because the cued information is already selected and ready to be acted on, cued probes result in faster reaction times compared to non-cued and new probes. Consistently, decreases in alpha power have been previously associated with increased neural firing^9^ and improved behavioral performance^9,29,30^, suggesting that the observed target-related alpha power decrease reflects a signature of target prioritization, that allows for a fast adaption to current task demands. More specifically, it has been hypothesized that such a prioritization may result from the attentional strengthening of relevant items to their (spatial) context^14^.

Whether in addition to a prioritization mechanism, active inhibition is also involved in the attentional selection from working memory remains a matter of debate. The majority of previous studies, using completely lateralized stimulus displays, do not allow for a clear distinction between an inhibition or a prioritization account^31^. Here, we demonstrated a distractor-related alpha power increase when participants were cued to re-orient their attention to a non-lateralized (i.e., central) working memory item, or – in other words – when shifting their attention away from a lateralized distractor. Increases in alpha power have been linked to decreased neuronal firing rates^9^ and have been shown to occur over task-irrelevant cortical areas^32^ as well as in anticipation of distractors^33^. Hence, interpreting the observed alpha power increase in terms of an inhibition mechanism appears plausible. Yet, the exact nature of such an inhibitory attentional signature still remains elusive. That is, if we conceptualize distractor inhibition as a top-down control mechanism that deteriorates the respective representations^34^, the question brought forth above likewise applies (cf. 3.1): If non-cued items are actively inhibited, why do non-cued items still cause interference when presented as a probe stimulus? Accordingly, both behavioral as well as neural evidence question the notion that an active inhibition mechanism completely deteriorates the non-cued item: On a behavioral level, using a series of cues, it has been shown that originally cued and then defocused items remain strengthened and result in faster and more accurate performance compared to trials in which a previously un-cued item (and thus “un-strengthened” item) is probed. In addition, EEG studies cast doubt on whether alpha lateralization actually mediates potential distractor exclusion effects^35,36^: For instance, Noonan and colleagues failed to find a lateralization of alpha power in response to a pre-cue indicating the location of an upcoming distractor, although, on the behavioral level, clear reaction time and accuracy improvements provide evidence for effective distractor exclusion (see also de Vries et al.^37^). However, it should be considered that the latter findings refer to anticipatory suppression of distractors in the visual domain, whereas the present study deals with the inhibition of previously relevant information within auditory working memory that subsequently becomes irrelevant.

Taken together, it remains possible that alpha enhancement contralateral to the distractor may be related to the temporal suppression of maintenance of information in a state of persistent activity (i.e., within the focus of attention), which does not necessarily alter the quality of unattended working memory contents^22,38^. Alternatively, we could conceptualize distractor inhibition as a mechanism that ultimately aims to permanently remove irrelevant items from working memory. Such a permanent removal has been conceptualized to operate through an un-binding mechanism, that unties the present item-context bindings^39^, rather than affecting the representation of the item itself. This is in line with recent evidence, demonstrating that alpha oscillations carry information about the maintained location (i.e., the “context” in the present case) but *not* about location-unrelated stimulus-features (i.e., the sound’s identity)^25^. Given that the present behavioral results argue against a complete removal of the un-cued item, such a mechanism would be expected to unfold gradually, resulting in weakened but not entirely removed representations^40^.

### 3.3 A dual mechanism underlying alpha lateralization

Overall, although we can only speculate on the exact nature of an inhibitory attentional control mechanism within auditory working memory, it is clear that we can dissociate distinct target- and distractor-related modulations of posterior alpha power. This supports the notion that working memory is a highly-flexible system that allows for the dynamic adaption to changing task demands by means of a dual-control mechanism. Alpha oscillations may serve as the underlying substrate that allows for flexible control over the state of current working memory representations, switching between prioritized and deprioritized representations depending on their current relevance for the task^36^. That the flexible prioritization and de-prioritization of working memory representation may, in fact, be mediated by two independent mechanisms, has been supported by a number of recent studies: Schneider et al.^6^ independently manipulated the meaning of a retro-cue such that it either indicated the to-be-remembered (remember-cue) or the to-be-forgotten (forget-cue) item(s). Their results indicated that both types of retro-cues elicited a target-related decrease in alpha power as well as a distractor-related increase in alpha power. Interestingly, the “cued process” (i.e., inhibition in forget-cue trials and target prioritization in remember-cue trials) always emerged first and was only later followed by its complementary counterpart (see also Poch et al.^41^). In addition to those studies revealing latency effects, Wöstmann and colleagues^42^ were able to demonstrate that alpha lateralization for target selection and distractor suppression were both different in strength and source origin as well as statistically uncorrelated. Evidence for the assumption that the strengthening of target representations can in fact occur independently from the de-prioritization of irrelevant information also comes from behavioral findings, illustrating how strategic considerations may affect the underlying control mechanisms^43^: When a retro-cue indicated the to-be-probed item with low validity (i.e. valid cue in 50% of trials), invalid-cue costs were largely absent, while participants still showed a clear benefit for validly cued trials. In line with the strengthening hypothesis, such a pattern of results suggests that it is, in fact, possible to prioritize or strengthen the relevant information without affecting the non-cued, irrelevant information. However, under conditions of high cue validity (i.e., valid cue in 80% of trials), which are more comparable to the present study, high invalid-cueing costs were observed, suggesting that in addition to a target prioritization mechanism, non-cued items were “removed” in order to free working memory resources.

### 3.4 Alpha lateralization is absent when spatial information becomes irrelevant

We did not observe any lateralization of alpha power following a neutral retro-cue, in which case both items maintained in working memory remained relevant. This appears plausible, considering that the neutral retro-cue did not require a re-orienting of spatial attention. In addition, this is consistent with the notion that auditory alpha lateralization is limited to situations in which spatial information is task-relevant^8,44^. Accordingly, once the neutral retro-cue appeared, the spatial location of the sounds became irrelevant to solving the task, as it was sufficient to know the sounds’ identity. In turn, this also means that alpha lateralization in response to a selective retro-cue reflects the spatially-specific access to the contents of working memory. This further corroborates the above-mentioned claim that alpha suppression indicates the strengthening of item-context bindings in working memory (cf. 3.2). In the present case, spatial location represents the item’s context, which is required to successfully distinguish cued from non-cued items. In contrast, in neutral retro-cue trials, knowledge of the spatial context is not required to solve the task. Those findings clearly emphasizes a critical difference to the visual domain, for which it has been shown that alpha-band activity tracks the stimulus position within working memory, irrespective of its relevance for the task.^27^

### 3.5 Bilateral alpha power suppression reflecting processing demands

In addition to hemispheric asymmetries in posterior alpha power, the present study also revealed a bilateral decrease following the presentation of the retro-cue, which was significantly more pronounced in distractor lateral trials compared to target lateral and neutral trials (cf. Fig. 3 f-h). Desynchronized alpha activity, resulting in low levels of alpha power (i.e., small amplitudes), has been associated with states of high excitability^9^ and is commonly interpreted as a mechanism reflecting functional engagement and information processing^45^. Accordingly, the event-related desynchronization of alpha power has been associated with stimulus processing (as opposed to ‘idling’)^46^, increased memory load^10^, and greater semantic elaboration^47^. Related evidence for alpha suppression as a general signature of information processing demands comes from the long-term memory literature: A recent study showed that on a single trial level, a greater decrease in alpha power was negatively correlated with a measure of stimulus-specific information represented in the BOLD signal^48^. The authors propose that alpha power decreases as a proxy for information processing; that is, greater decreases indicating greater information processing demands. In line with this interpretation, distractor lateral trials in the current study presented the acoustically most challenging spatial condition, because the to-be-attended (central) sound was originally embedded by two-neighboring sounds. Thus, one may speculate that the representation generated at encoding is likely to be of lower quality than that of the lateral sound stimuli and may thus require more attentional resources to be re-focused within working memory. Griffith and colleagues emphasize that such an alpha power decrease does not carry information about the stimulus itself, but does in contrast provide conditions under which the representations of a required stimulus are optimally recalled. It is worth noting that there is a number of studies showing contradicting patterns of results: For instance, alpha power has been shown to increase with higher acoustic detail in a speech distractor^49^ or as a function of memory load^50^. Those findings are typically interpreted in terms of an inhibition account. However, in the present study, inhibition is not expected to be reflected by overall, non-lateralized alpha power, as the to-be-inhibited item was lateralized.

## 4. Conclusion

Taken together, using a retro-active cueing design, we demonstrate that it is possible to unambiguously dissociate target- and distractor related modulations of alpha power oscillations. The results indicate that both excitatory and inhibitory attentional control mechanisms contribute to the selection of information from working memory. Accordingly, shifts of attention toward a lateralized target resulted in a contralateral decrease in alpha power, whereas shifting attention away from a lateralized distractor resulted in a contralateral increase in alpha power. The pattern of results support the notion that alpha lateralization mediates the task-dependent prioritization and de-prioritization of items within working memory^36^. In addition, we strengthen the previous claim that auditory alpha lateralization is absent when spatial information is task-irrelevant^8,44^.

## 5. Methods

### 5.1 Participants

Twenty-eight volunteers were paid to participate in this study. Four participants had to be excluded due to technical problems with their EEG recordings. Four additional participants showed excessive eye-movement artefacts (containing excessive lateral eye movements on more than 1/3^rd^ of all trials) and were therefore excluded from further EEG analyses. The remaining twenty participants (12 female) were aged between 19 and 28 years (mean age: 23.4 years) and right-handed as assessed using the Edinburgh Handedness Inventory^51^. All participants reported normal or normal-to corrected vision, no history of or current neurological or psychiatric disorders. Hearing acuity was assessed using an audiometry, including eleven pure-tone frequencies (0.125 – 8 kHz; Oscilla USB 100, Inmedico, Lystrup, Denmark). Hearing thresholds indicated normal hearing (≤ 25 dB hearing level). The study was conducted in accordance with the Declaration of Helsinki and approved by the Ethical Committee of the Leibniz Research Centre for Working Environment and Human Factors. Written informed consent was given by all participants prior to the beginning of the experimental procedure.

### 5.2 Experimental setup and stimuli

The experiment took place in a dimly lit, echo-attenuated, sound-proof room. Participants were seated in front of a semi-circular array of eight broad-band loudspeakers (SC5.9; Visaton, Haan, Germany; hoursing volume 340 cm^3^) mounted in the horizontal plane. Three of those loudspeakers, located at azimuthal positions of −90°, 0°, 90°, were used for sound presentation in the present study. One additional loudspeaker was located right behind the participant’s head at a distance of approximately 30 cm. A red light-emitting diode, attached below the central loudspeaker (diameter 3 mm, turned off) served as a central fixation point. The participants’ head was kept at a constant position using a chin rest.

Eight familiar animal sounds, chosen from an online sound archive^52^, served as experimental stimuli. The original sounds (‘birds chirping’, ‘dog barking’, frog broaking’, ‘sheep baaing’, ‘cat meowing’, ‘duck quacking’, ‘cow mooing’, ‘rooster crowing’) were cut to a constant duration of 600 ms (10 ms on/off ramp), while leaving the spectro-temporal characteristics unchanged. In addition, syllable sounds served as cue-stimuli, indicating a certain subset of sound positions as relevant in a given trial. That is, the first two or three letters of the German words for right (i.e. “rechts”), left (i.e. “links”), middle (i.e. “mitte”), and both (i.e. “beide”) were used to construct the six cue-stimuli (“*li*”, “*re*”, “*mi*”, “bei”, “*mi-li*”, “*mi-re*”). The words were spoken by a female speaker (mean pitch 199 Hz). All cues had a duration of 400 ms. The overall sound pressure level of the sound arrays (presented at frontal loud speakers) was about 65 dB(A), whereas single animal sounds and the speech cue-stimuli (presented from behind participants’ heads) were presented at a sound pressure level of 60 dB(A).

### 5.3 Procedure and task

The present experiment is a modified and simplified version of a retroactive cueing paradigm previously used in an investigation of visual retroactive attentional orienting^6^. Each trial comprised a sequence of four acoustic stimulus events, consisting of pre-cue, sound array, retro-cue, and probe. A trial started with the presentation of a pre-cue (400 ms), instructing participants to attend either the central and the left loudspeaker (“*mi-li*”) or the central and the right loudspeaker (“*mi-re*”). Thus, in the subsequently upcoming sound array, containing three animal vocalizations, one lateral sound was always known to be completely irrelevant for the rest of the trial. The sound array appeared 1000 ms after the pre-cue and was presented for 600ms. Following a short delay (1000 ms), a retroactive cue (400 ms) indicated either one (i.e., selective retro-cue) or both (i.e., neutral retro-cue; “*bei*”) of the items currently maintained in working memory as further relevant for the task. Selective retro-cue trials can be further subdivided into target lateral trials (i.e., the retro-cue indicated the lateral item as further relevant, “*li*” or “*re*”) or distractor lateral trials (i.e., the retro-cue indicated the central item as relevant whereas the lateral item becomes irrelevant, “*mi*”). Each of the four retro-cue types was presented equally often (25% of trials), to enforce that none of the sound positions were favored irrespective of cue-type. That resulted in a higher number of trials for target lateral retro-cues (“li” and “re”; 50% of trials) as opposed to distractor lateral (“mi”, 25% of trials) and neutral (“bei”, 25% of trials) retro-cues. Finally, 1000 ms subsequent to the retro-cue, a probe stimulus was presented. The probe stimulus was either the item that became irrelevant following the retro-cue (“non-cued probe”), the retro-cued item (“cued probe”), or a sound stimulus that never appeared in a given trial before (“new probe). Note that the animal vocalization which was initially presented at the third, irrelevant position in the sound array (as indicated by the pre-cue) never appeared as a probe to ensure that participants would always ignore it. Because non-cued and new probes required a NO response they constituted 25% of all trials, respectively, whereas cued probes, requiring a YES response, constituted 50% of all trials. Participants were instructed to indicate whether the probe item matched (one of) the retro-cued item(s) by giving a YES (50% of trials) vs. NO (50% of trials) response. Participants responded by pressing one out of two vertically aligned keys, using the index finger and thumb of their right hand. The assignment of response alternatives (yes vs. no) to keys was counterbalanced across participants. Responses had to be given within 1500 ms following probe offset. All probe stimuli, as well as pre- and retro-cues, were presented from a loudspeaker located in the median plane behind the participant’s head. The experiment consistent of a total of 30 practice trials and 800 experimental trials. The latter were divided into eight task blocks of 100 trials each. In-between blocks, short self-paced breaks served the prevention of fatigue in the course of the experiment. All participants were presented with the same, pseudo-random sequence of trials.

### 5.4 Data analysis

#### 5.4.1 Behavioural data

Reaction times and percentage of correct responses served as measures of behavioural performance. Trials with responses that occurred after the pre-defined response period (i.e., within the inter-trial interval) were considered invalid. For analysis of reaction times, only correctly answered trials were included. In order to verify that the paradigm resulted in a retro-cue effect ^18^, we performed a paired sample *t*-test and a Wilcoxon signed-rank test, contrasting selective versus neutral retro-cues, for reaction times and accuracy data, respectively. Note, that for this comparison the non-cued probe trials (which were not present for neutral retro-cues) were excluded to avoid an imbalance between conditions. In addition, to test for the hypothesized interference effect in non-cued probe types, we contrasted the different probe types within the selective retro-cue condition by means of a repeated-measures analysis of variance (ANOVA) including the factor *probe type*. Again, the ANOVA was run for both reaction times and accuracy. Since for both the reaction time data as well as the accuracy data, we conducted a total of two analyses on the same data set (i.e., test for retro-cue type as well as for probe type), *p*-values were fdr-corrected for multiple comparisons across those two sets of analyses (cf. 5.4.5). In addition, post-hoc paired-sample *t*-tests (or their non-parametric alternative, i.e., Wilcoxon signed-rank test) were conducted and corrected for multiple comparisons. Note that adjusted *p*-values can be greater than 1.

#### 5.4.2 EEG recording and processing

The EEG was recorded with a sampling rate of 1000 Hz from 64 Ag/AgCl passive electrodes (Easycap GmbH, Herrsching, Germany), using a QuickAmp-72 amplifier (Brain products, Gilching, Germany). The electrodes were arranged across the scalp according to the extended 10/20 scalp configuration. AFz served as the ground electrode. The average of all channels constituted the online-reference. Impedances were kept below 10 kΩ during recording. Further pre-processing of the data was run in MATLAB® (R2018b) and EEGLAB^53^. First, a Hamming windowed sinc FIR high-pass (0.5 Hz; 0.25 Hz cut-off frequency, 0.5 Hz transition band width, filter order: 6600) and low-pass filter (30Hz; 30.25 Hz cut-off frequency, 0.5 Hz transition band width, filter order 440) were applied. Then, channels with a normalized kurtosis (20% trimming before normalization) exceeding 5 standard deviations of the mean were rejected, using the automated channel rejection procedure implemented in EEGLAB (M = 3.25 channels, range = 1 – 5). Anterior lateral channels (Fp1/2, AF7/8, AF3/4, F9/10) were excluded from channel rejection to ensure reliable identification of eye movements. Data were re-referenced to the average of all non-rejected channels. Epochs ranging from −1000 to 7500 ms relative to the onset of the pre-cue were generated. A rank-reduced independent component analysis (ICA) was run on a subset of the data, down-sampled to 200 Hz and including every second epoch. To detect and remove independent components (ICs) reflecting eye blinks, vertical eye movements, and generic discontinuities, the EEGLAB plugin ADJUST^54^ was applied. In addition, for each IC, a single-equivalent current dipole model was estimated by means of a spherical four-shell head model^55^. Any components with a dipole solution exceeding a threshold of 40% residual variance were also removed. Taken together, on average 20.75 ICs (range = 7 – 33) were rejected. This was followed by an automatic trial rejection procedure, detecting and removing data epochs, containing data values exceeding a threshold of 5 standard deviations in an iterative procedure (threshold limit: 1000µV, maximum % of trials rejected per iteration: 5%). On average, 163.3 (20.41%) trials (range 0 - 309) were rejected in the course of this procedure. Finally, data from channels that were originally rejected were replaced using spherical interpolation. For faster processing of time-frequency analyses (cf. below), the data were down-sampled to 500 Hz.

#### 5.4.3 Time-frequency analysis: Alpha lateralization

In order to extract spectral power, the epoched EEG data was convoluted with a complex Morlet wavelet. The frequencies of the wavelets ranged from 4 Hz to 30 Hz, increasing logarithmically in 52 steps. Convolution began with a 3-cycle wavelet for the lowest frequency and increased linearly as a function of frequency with a factor of 0.5, resulting in a 11.25-cycle wavelet for the highest frequency. No spectral baseline-correction was applied, since the calculation of alpha lateralization indices requires raw power input. The resulting event-related spectral perturbation (ERSP) epochs ranged from −582 to 7080 ms relative to pre-cue onset. Lateralized effects in posterior alpha band power (8-13 Hz) in response to the retro-cue were assessed as a measure of retroactive attentional orienting. As a robust measure of lateralization, a lateralization index^56^ was calculated as follows:

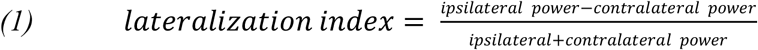

Mean power across trials was extracted for each subject ipsilateral and contralateral relative to the lateralized item in a given trial (i.e. the target in target lateral and neutral trials or the distractor in distractor lateral trials). Based on the electrode cluster used by Schneider et al.^6^, whose design this study was based on, power was averaged across the following electrodes of interest: PO7/8, P7/8, P5/6, and PO3/4. Because we had a clear hypothesis regarding an expected difference between the target lateral and the distractor lateral conditions, we first contrasted those two conditions using a cluster-based permutation approach: In a first step, the original data from the two conditions were contrasted by means of paired *t*-tests, comparing the lateralization index at each time-point (i.e., 200) and each frequency (i.e., 52). Note that no space dimension exists here, because power was already averaged across the electrodes mentioned above. This resulted in a time by frequency matrix of *p*-values. In a second step, the condition labels (i.e., target lateral vs. distractor lateral) were randomly assigned to the actual data points and again, a paired *t*-test was run for each time-frequency point. This procedure was iterated 1000 times and thus, resulted in a 52 x 200 x 1000 matrix of *p*-values. For each iteration, the size of the largest cluster of time-frequency points with a *p*-value below .05 was determined, resulting in a distribution of maximum cluster sizes to be expected under the null hypothesis. The 95^th^ percentile of this distribution served as a cut-off value against which the clusters from the true data were compared. That is, only clusters of time-frequency points with a *p*-value below .05 that were larger than the 95^th^ percentile of the permutation-based distribution of maximum cluster sizes were considered significant.

Based on the approximate cluster time-window in the alpha frequency range (8-13 Hz) derived from this comparison, we compared the neutral retro-cue condition to the target lateral as well as the distractor lateral condition by means of paired-sample *t*-tests. Note that time-windows determined based on a cluster-permutation approach should not be interpreted in term of an exact on- or offset of an effect^57^. In addition, we tested for significant alpha lateralization within conditions (i.e., one-sample *t*-tests against zero). *P* values were corrected for multiple comparisons (cf. *5.4.5*) including all five post-hoc comparisons (i.e., 3 one sample-*t*-tests within conditions and 2 paired sample *t*-test comparing conditions).

#### 5.4.4 Time-frequency analysis: Bilateral alpha desynchronization

For the analysis of bilateral alpha power desynchronization, baseline-corrected ERSPs were computed. Morlet wavelet convolution was conducted as described above (cf. 4.4.3.), but a spectral baseline was extracted for each frequency (−300 to 0 relative to pre-cue onset). Mean alpha power (8-13 Hz at PO7/8, P7/8, P5/6, and PO3/4) was extracted for each subject and the three conditions (i.e., target lateral, distractor lateral, neutral), respectively. To test for overall differences in alpha suppression between conditions, a one-way repeated measures ANOVA was performed. Paired sample *t*-tests were conducted for post-hoc comparisons. The resulting *p*-values were corrected for multiple comparisons (cf. *5.4.5*).

#### 5.4.5 Inferential statistics and effect sizes

All statistical analyses of the data were conducted in MATLAB® (R2018b). Behavioral data were formatted and prepared for analysis in MATLAB using RStudio (Version 1.2.1335). Parametric tests were applied when the data met the normality assumption (*p* > .05 in Lilliefors test). Otherwise, appropriate non-parametric tests, including Friedman’s test and Wilcoxon signed rank test, were conducted. Mauchly’s test for sphericity was performed for all repeated-measures ANOVA models. In case of significant violations of sphericity (*p* <.05), Greenhouse Geisser correction was applied. The significance of effects was assessed at an alpha level of .05. Reported *p*-values associated with tests based on the *F*-distribution are directional, given that the *F*-distribution is not symmetrical. All conducted (non-)parametric *t*-tests were two-tailed. As measures of effect size, partial eta-squared (*ηp*^2^) is provided for repeated-measures ANOVA. Hedges’s *g* and *g*_1_ served as an effect size measure for paired and one-sample *t*-tests, respectively. Cohen’s U3 (in the following denoted as U3) is provided for non-parametric Wilcoxon signed rank tests. The latter can be interpreted as a measure of overlap between two distributions, ranging from 0 to 1, where 0.5 indicates no effect. Hedges’s *g*, *g*1, and U3 are calculated using the MATLAB Toolbox ‘Measures of Effect Size’^58^. To correct for multiple comparisons, false discovery rate (FDR) correction was applied^59^. Corrected *p*-values are denoted as *p*_adj_.

To further facilitate the interpretation of results, we additionally report the Bayes factor (BF). While classical null hypothesis significance testing only allows conclusions on whether we can disprove the null hypothesis, the BF also allows for an assessment of whether the data favors the null hypothesis compared to an alternative hypothesis (see ^60,61^ for a general introduction to Bayesian hypothesis testing). A BF greater than 3, 10, 30, and 100 provides moderate, strong, very strong, and extreme support for the alternative hypothesis^60^. Values in-between 0.33 and 3 are usually interpreted as anecdotal evidence, whereas values lower than 0.33, 0.1, 0.03, and 0.01 indicate moderate, strong, very strong, and extreme evidence in favor of the null hypothesis^60^. BFs were calculated using the MATLAB BF functions and default priors implemented by Krekelberg^62^.

## 6. Data availability

The datasets generated in the course of present study are stored in the Leibniz Research Centre repository and are available from the corresponding author upon request.

## 8. Acknowledgements

The authors would like to thank Kimberly Freytag and Stefan Weber for their contribution to data collection.

## 9. Author Contributions

L.I.K., D.S. and S.G. designed the study. Data collection was performed by L.I.K. and A.B. L.I.K. performed the data analysis and interpretation under supervision of D.S. and S.G. L.I.K. drafted the manuscript. All authors contributed to the revision of the manuscript and approved the final version of the manuscript for submissions.

## 10. Additional Information

### Supplementary material

Supplementary material associated with this article can be found in the online version.

## Appendix A. Supplementary Material

### 1. Control analysis: Lateral eye movements prior to retro-cue presentation

Previous related experiments from the visual domain raised concerns about alpha power asymmetries being potentially confounded by lateral shifts in gaze position^1,2^. Figure S1 shows that the frontal event-related potentials (ERP) contain a minor fixation offset towards the lateralized item in the sound array prior to retro-cue onset. Since gaze position has been shown to affect the perceived sound eccentricity^3^, lateral saccadic eye movements may have likewise affected the electrophysiological correlates of attentional orienting towards perceived sound locations. We performed several control analyses to rule out such contamination of our data:

First of all, we calculated correlations between single-trial indices of lateral saccadic eye movements and posterior alpha power asymmetries. The ipsilateral minus contralateral difference in ERP amplitude at fronto-lateral channels F9/10 across the 200 ms interval prior to retro-cue onset served as a measure of single-trial lateral saccadic eye movements. That is, for left-sided targets or distractors (depending on condition), ERP amplitudes at F9 minus F10 were subtracted, whereas for right-sided targets and distractors ERP amplitudes at F10 minus F9 were subtracted. Note that electrode positions F9 and F10 correspond to the most frontal channels in our EEG setup and are thus comparable to typical hEOG channel locations. Hemispheric asymmetries in the posterior alpha frequency band were analogously computed by calculating the lateralization index, as described in the methods section of the manuscript (cf. 5.4.3). On a single-trial level, this was done separately for right-sided and left-sided targets/distractor. Single-trial alpha asymmetries were assessed in the same frequency range, time interval and electrode cluster used for the main analysis (i.e., 8-13 Hz, electrode cluster: PO7/8, P7/8, P5/6, PO3/4, time window: 700 – 1300 ms post retro-cue onset). Pairwise Spearman’s Rho or Pearson correlation coefficients (depending on normality properties of the data) were then calculated for each subject and each condition (i.e., target left, target right, distractor left, distractor right). After Fisher-Z transforming the correlation coefficients, one-sample *t*-tests were conducted in order to test for a statistically reliable relation between lateral eye movements (prior to retro-cue onset) and alpha lateralization (following the retro-cue) in each condition. The resulting *p*-values were FDR-corrected for multiple comparisons. The scatter plots in figures S2 and S3 illustrate that there was no apparent relationship between the two measures. The analysis confirmed this, revealing that the single-subject correlation coefficients were not significantly different from zero, neither for target lateral nor for distractor lateral trials (all *t* < .01, *p* > .9, *p*_adj_ < 2.46, BFs < .24). Not that we did not perform this control analysis for neutral trials, since we did not observe any significant lateralization of alpha power in that condition.

In addition, since the above-mentioned correlative approach relies on the presence of null findings (i.e., a non-significant correlation between lateral saccadic eye movements and alpha lateralization), we also ran a repeated-measures analysis of covariance, including the within-subject factor *retro-cue type* (distractor lateral vs. target lateral) and as a covariate, the ipsilateral minus contralateral portions of the grand average ERP signal (relative to the lateral target or distractor position) at electrodes F9/10 across both conditions. The latter did not include the neutral retro-cue condition. The main effect of *retro-cue type* remained significant (*F*_(1,18)_ = 12.84, *p* = .002, *ηp*^2^ = 0.42), while there was no significant interaction of *retro-cue type* and *saccades (F*_(1,18)_ = 3.28, *p* = .09, *ηp*^2^ = 0.15). Taken together, these analyses argue against any confounding influence of lateral eye movement patterns on posterior alpha lateralization.

---

**Supplementary figure S1.**
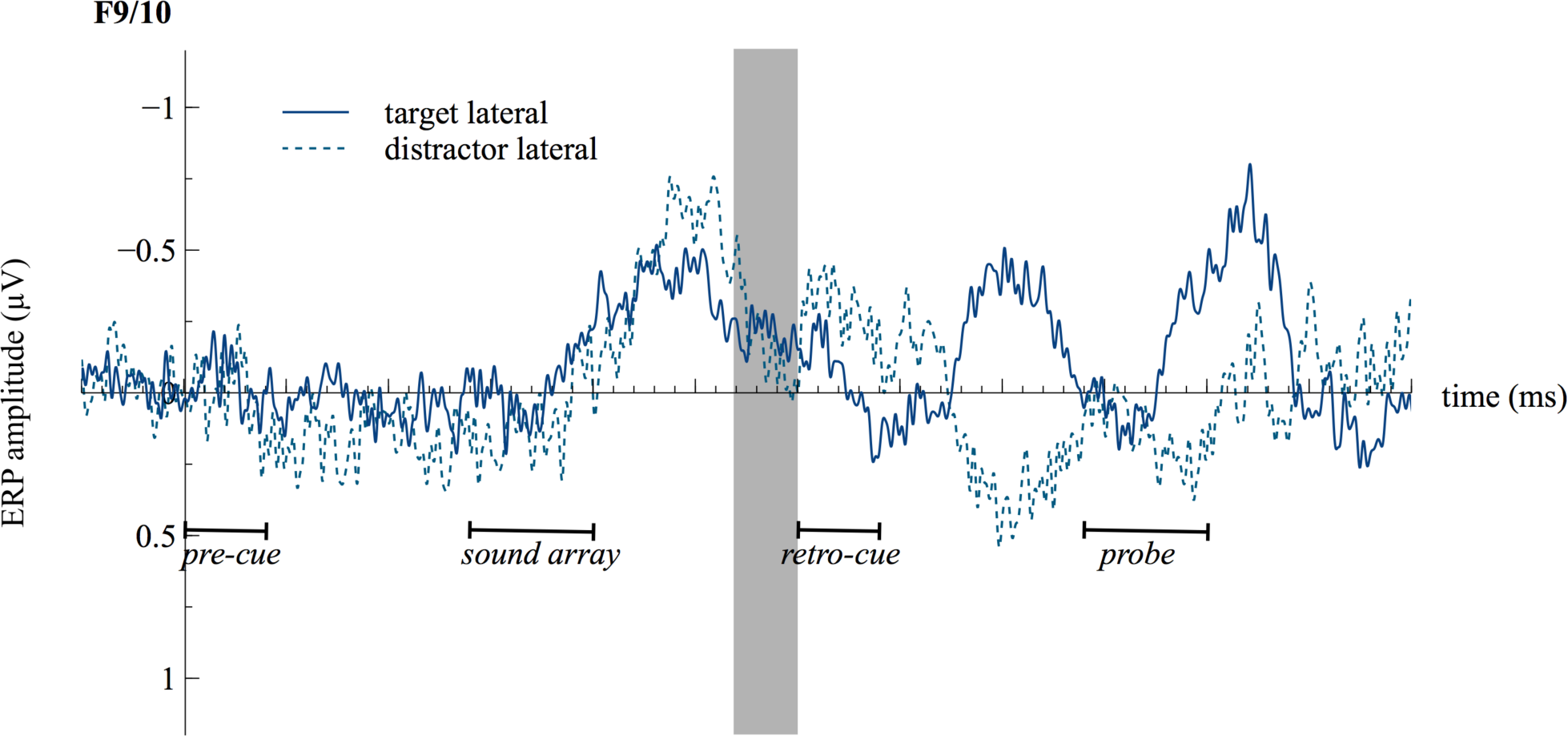
Contralateral minus ipsilateral event-related potentials (ERPs) at fronto-lateral channels F9/F10. The grey area marks the time interval used to extract single-trial ERP amplitudes as a measure of lateral saccadic eye movements. Note that for reasons of consistency with the Alpha Lateralization Index (ALI), the analysis was performed using the ipsilateral minus contralateral amplitude differences.

**Supplementary figure S2.**
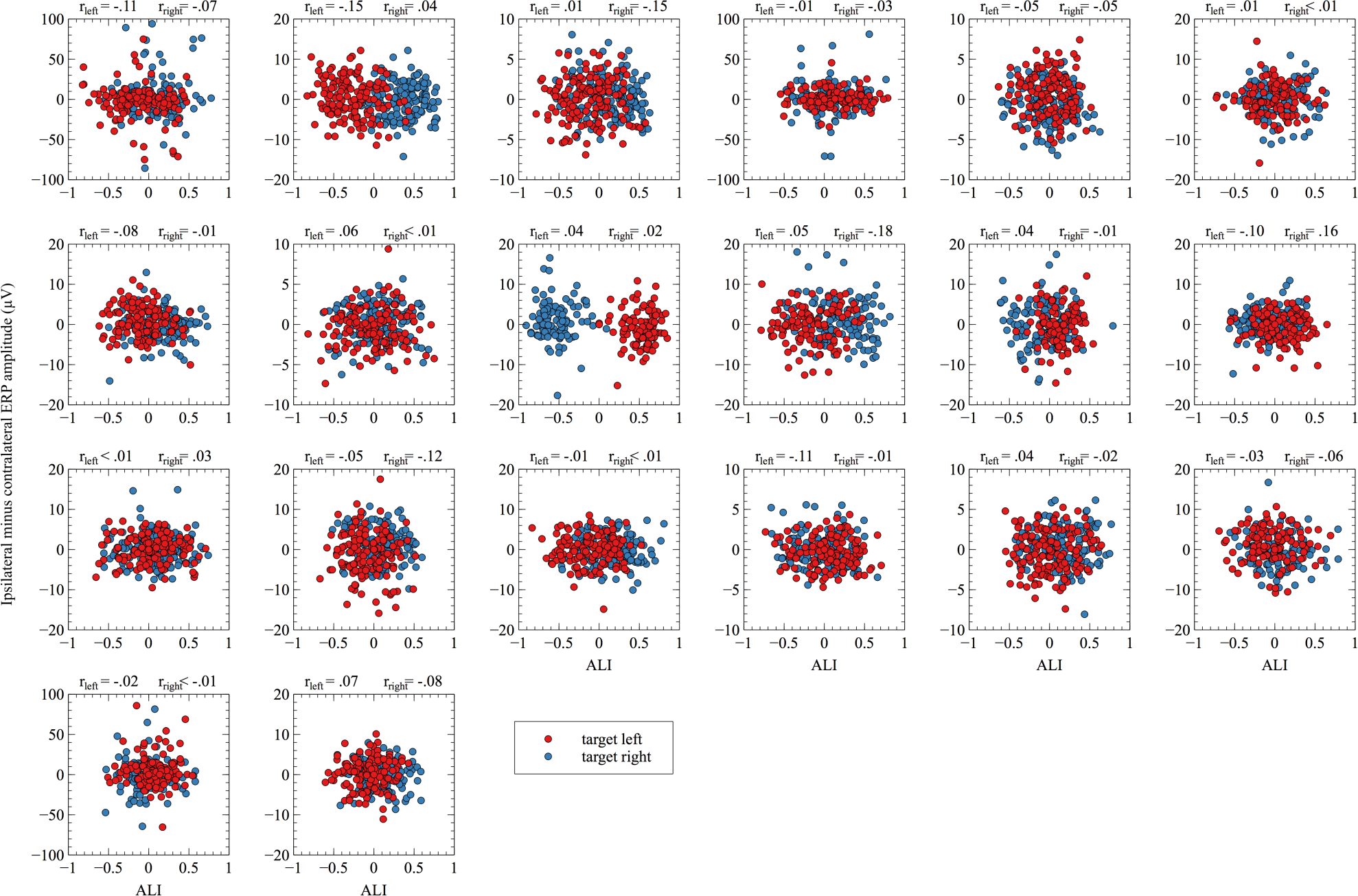
Scatter plots illustrate the association between posterior alpha asymmetries and lateral saccadic eye movements in target lateral trials. Each plot corresponds to the data of a single subject.

**Supplementary figure S3.**
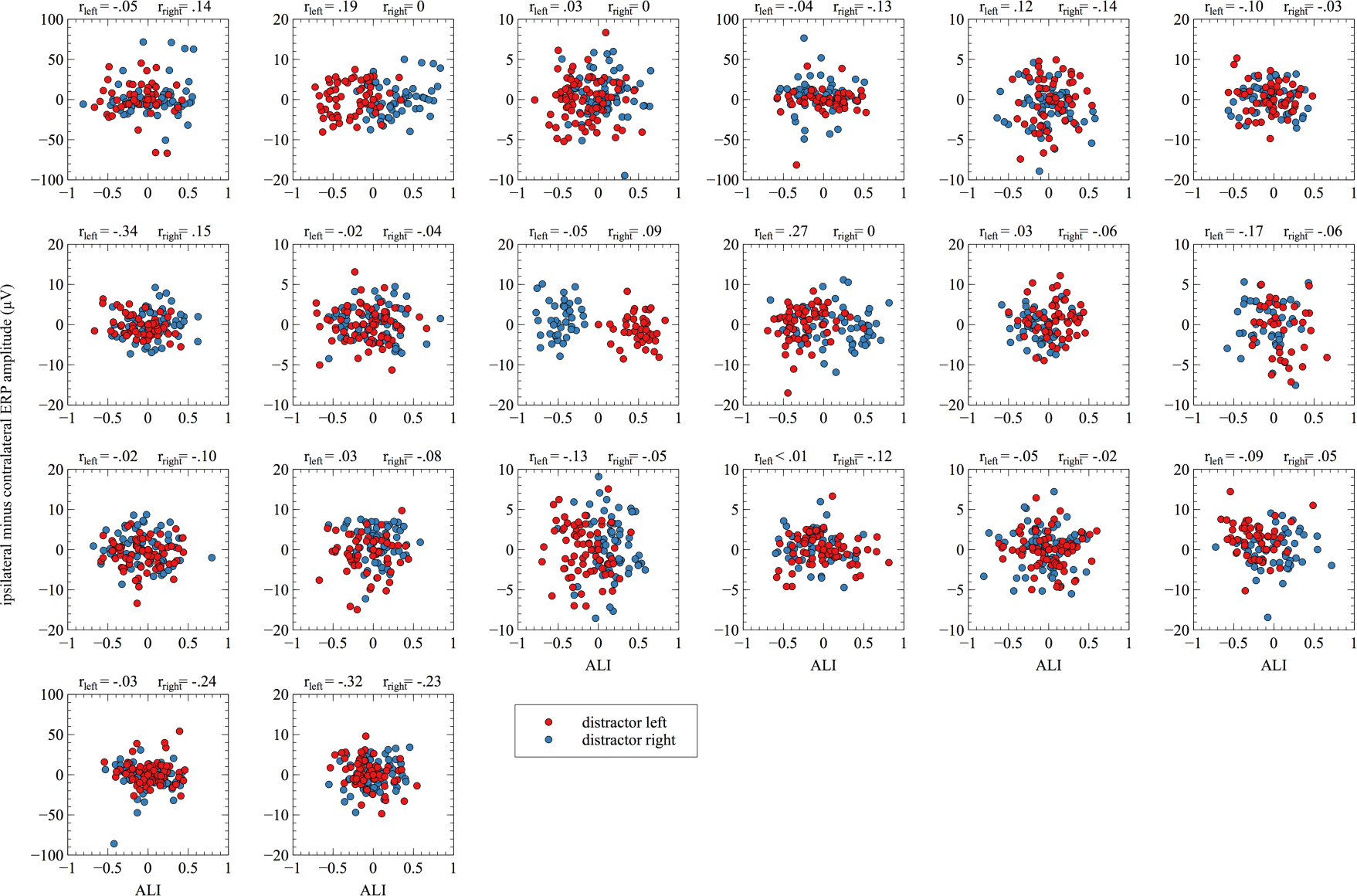
Scatter plots illustrate the association between posterior alpha asymmetries and lateral saccadic eye movements in distractor lateral trials. Each plot corresponds to the data of a single subject.

